# Evaluation of solvents used for fabrication of microphysiological systems

**DOI:** 10.1101/2021.03.24.436761

**Authors:** Xiaopeng Wen, Seiichiro Takahashi, Kenji Hatakeyama, Ken-ichiro Kamei

## Abstract

Microphysiological systems (MPSs) have shown great promise for the advancement of drug discovery and toxicological tests, and as an alternative to animal models. However, although several chips and systems have been reported, some important issues are yet to be addressed, such as the use of polydimethylsiloxane (PDMS). Cyclo olefin polymers (COPs) have advantages over other thermoplastic materials, but most COP-based MPSs use solvent bonding during fabrication, which can affect any cells they are used to culture. This study uses a photobonding process with vacuum ultraviolet (UVU) to produce MPSs without the need for solvents such as cyclohexane, dichloromethane, and toluene. This is then used for comparison to investigate the effects of solvents on cell cultures. Quantitative immunofluorescent assays show that the coating efficiencies of extracellular matrix proteins, such as Matrigel and collagen I, are reduced on solvent-treated COP surfaces, compared with those prepared using VUV photobonding. Furthermore, SH-SY5Y neuroblastoma cells are used to evaluate cytotoxicity. This shows that solvent-MPSs induce apoptosis, but VUV-MPSs do not. These results provide insights into solvent bonding for MPS fabrication so that undesirable reactions can be avoided. Moreover, this work may be used to standardize MPS protocols and establish good manufacturing practices.

## 1. Introduction

Microphysiological systems (MPSs) or organs-on-a-chip (OoC) hold great promise for the advancement of drug screening and toxicological testing because they can be used to predict the efficacy or toxicity of drugs without the use of experimental animals [1–5]. In recent years, ethical concerns have driven a move away from animal testing, and MPSs with human cells represent an effective alternative.

Most MPSs are based on microfluidic technology, and they often use polydimethylsiloxane (PDMS) or plastic materials as structural components. Although PDMS has been used for microfluidic cell cultures because of its biocompatibility, gas permeability, and optical transparency, it has some widely recognized limitations including the absorption of hydrophobic molecules, leaching of uncross-linked monomers, and water evaporation [6–10], which might interfere with cellular behavior and drug responses in MPSs. In addition, the soft lithography process used to manufacture PDMS-based MPSs is not suitable for mass production because the elasticity of PSMS microfluidic structures makes them difficult to handle [11]. Furthermore, PDMS-based MPSs cannot be stored long-term because a gradual cross-linking process causes structural shrinkage [8]. Alternatives to PDMS, including plastic materials, such as polystyrene (PS), polymethyl methacrylate (PMMA), and cyclo olefin polymer (COP), have been used for general cell cultures for a long time, and some MPSs have also been used [10–14]. These materials can be mass produced using techniques such as metal molding, so they have great potential in the realization of commercial MPS products [14,15].

However, plastic materials still have some limitations, such as the bonding processes required to assemble two or more plastic structures. Previously, many cell cultures and MPSs with plastic materials have relied on double-sided tapes, glues, or solvents for bonding [16]. Among these processes, solvent bonding is most practical due to its relative simplicity. However, the additional materials have an undesirable effect on cell cultures and assays due to leakages and microstructure disruptions, which can affect the responses of cultured cells. To address this, solvents are eliminated using a vacuum to reduce their effects, but small amounts remain and they can still affect cultured cells. Therefore, an alternative bonding process, that does not require additional materials, is needed to eliminate these problems.

Photobonding using vacuum ultraviolet (VUV) light from an excimer lamp is an ideal way to achieve this. Unlike glue- and solvent-based bonding processes, photobonding does not disrupt the microfluidic structures. Previously, we reported that VUV photobonding could be used to fabricate COP-MPSs, and that this reduced apoptosis of cultured human induced pluripotent stem cells (hiPSCs) compared with COP-MPSs fabricated with solvent bonding [17]. Thus, evidence suggests that VUV photobonding would be beneficial for cell cultures and assays in MPSs, and that solvent bonding affects cultured cellular phenotypes. In other words, COP-MPSs with VUV photobonding make it possible to investigate the effects of solvent bonding on cell cultures and assays, which is known from experience, but the underlying mechanisms are not understood.

This study investigates the effect of solvent bonding during MPS fabrication on cell cultures and assays. For evaluation, typical solvents such as toluene, cyclohexane, and dichloromethane, and fabricated COP-MPSs are used. COP-MPSs fabricated with VUV photobonding (COP-MPS-V) are used for comparison. The effects of these MPSs on extra-cellular matrix (ECM) protein coatings, which are generally used to facilitate cell adhesion and growth, are evaluated. Finally, SH-SY5Y cells, which are widely used to assess the cytotoxicity of chemicals/materials [18,19], are used to evaluate the cytotoxic effects of solvent-based COP-MPSs. This investigation provides insights into the undesirable effects on cell cultures caused by MPSs, and will aid the development of precise cell cultures and assays.

## 2. Materials and Methods

### 2.1. Fabrication of the COP microfluidic structure

The COP-MPS comprises two COP layers (microfluidic and bottom layers) and eight microfluidic cell-culture channels (800 μm wide, 7500 μm long, and 250 μm high). Each channel had a medium inlet and outlet (2 and 1 mm in diameter, respectively), and a total volume of 13.2 μL [17]. Two different COP-MPS microfluidic structures (RC2 and RC3) with different heights (280 and 320 μm, respectively) were prepared for the solvent and VUV photobonding processes, respectively. The methods used to fabricate the COP-MPS included metal molding (**Figure S1**). Briefly, this involved using two metal molds to fabricate each microfluidic structure. The COP material (Zeonex 480R, Zeon, Tokyo, Japan) was injected into the two molds individually, then removed from the molds to give the COP microfluidic structures.

### 2.2. Solvent-bonding process for COP-MPS fabrication

Solvent bonding was conducted with the same COP microfluidic structures (RC2) (**Figure 1A** and **Table 1**). Toluene, cyclohexane, and dichloromethane were obtained from FUJIFILM Wako Pure Chemical Corp. (Osaka, Japan). The bottom COP-MPS layer was exposed to solvent vapor, then the bottom COP-MPS and microfluidic layers were assembled using a hot press under the conditions listed in **Table 1**. After bonding, the remaining solvents were allowed to evaporate in a constant-temperature bath at 60 °C for 24 h.

**Figure 1.**
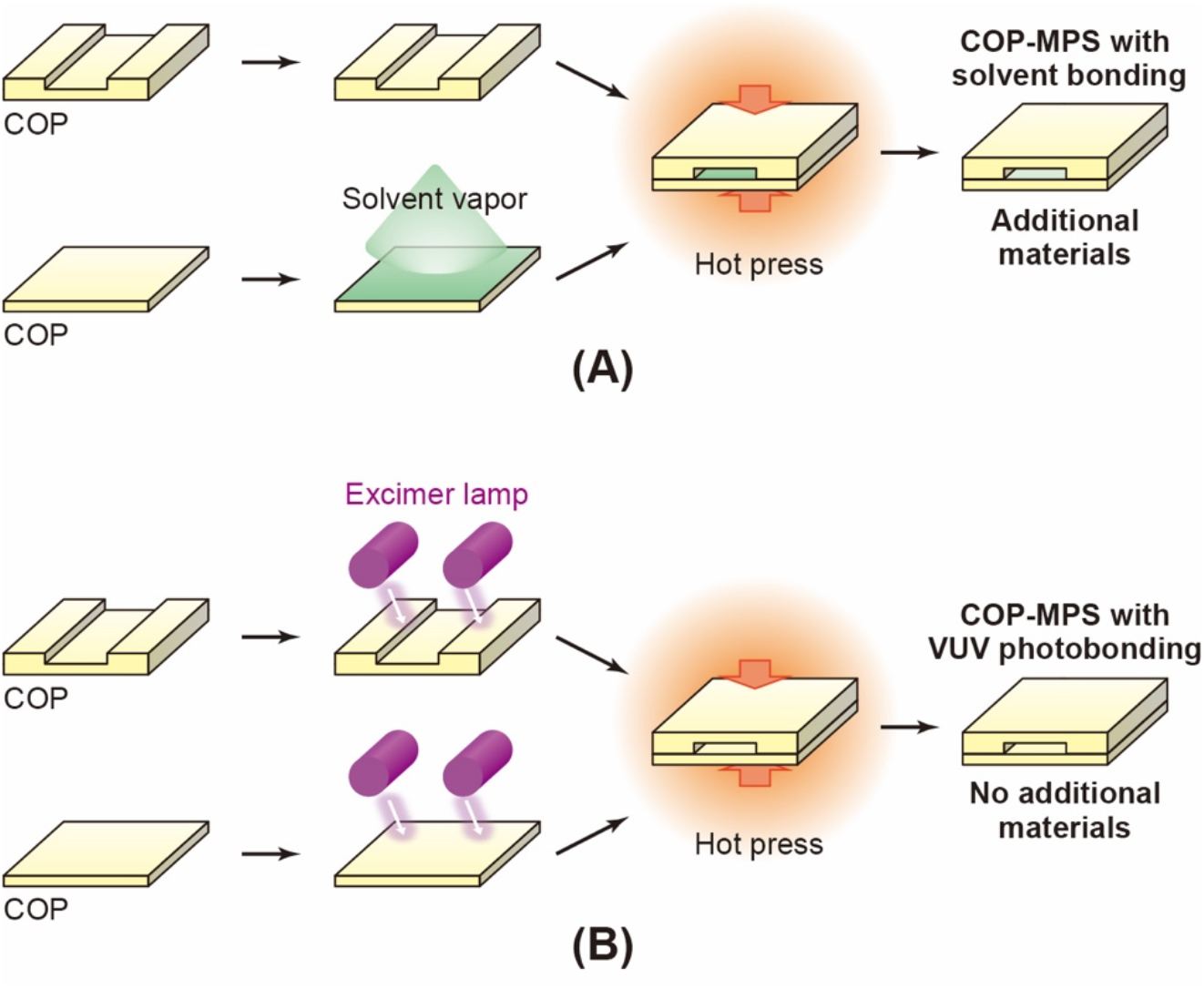
Schematic diagrams showing (A) solvent bonding and (B) VUV photobonding with an excimer lamp for the production of microphysiological systems made of cyclo olefin polymer (COP-MPS).

**Table 1.**
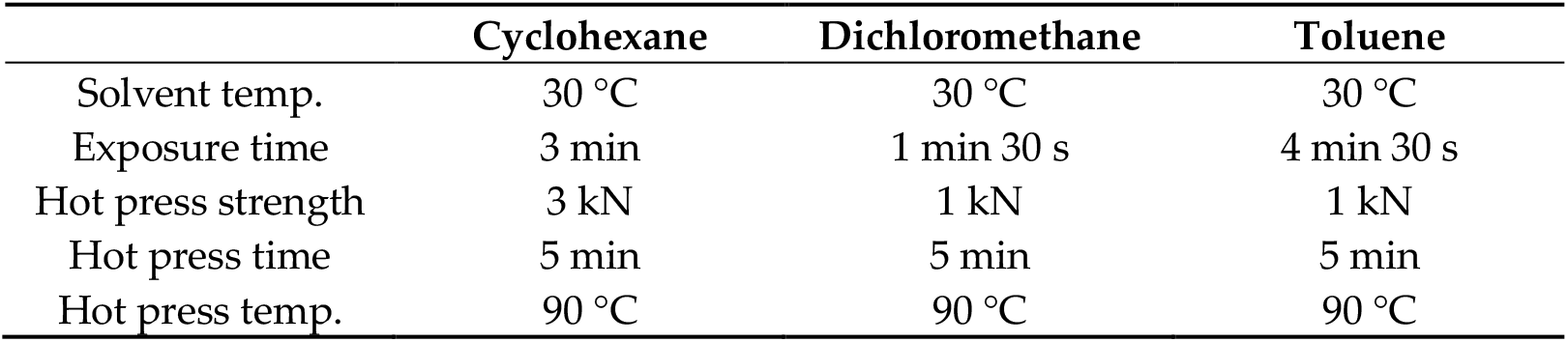
Parameters for the solvent-bonding processes.

|  | Cyclohexane | Dichloromethane | Toluene |
| --- | --- | --- | --- |
| Solvent temp. | 30 °C | 30 °C | 30 °C |
| Exposure time | 3 min | 1 min 30 s | 4 min 30 s |
| Hot press strength | 3 kN | 1 kN | 1 kN |
| Hot press time | 5 min | 5 min | 5 min |
| Hot press temp. | 90 °C | 90 °C | 90 °C |

### 2.3. VUV photobonding process for COP-MPS-V fabrication

Once the COP-MPS microfluidic structures (RC3) were removed from the molds, the components were irradiated with VUV using an excimer lamp source (172 nm; Ushio INC., Tokyo, Japan) at 25 °C. The surfaces of the components were then assembled using a heat press at < 132 °C (**Figure 1B**).

### 2.4. Coating COP-MPSs with extracellular matrices (ECMs)

To produce the ECM coatings on the COP-MPSs, 15 μL of 0.26, 0.52, and 5.15 mg/ml of Matrigel (Sigma-Aldrich, St. Louis, MO, USA), or 0.005%, 0.01%, and 0.1% (w/v) bovine collagen type I (Sigma-Aldrich, St. Louis, MO, USA) in Dulbecco’s modified Eagle medium (DMEM)/F12 (Sigma-Aldrich, St. Louis, MO, USA) was added to each microfluidic channel and kept at 4 °C for over 24 h. Excess Matrigel or collagen type I was removed, then the channels were washed with PBS.

### 2.5. Evaluation of the amount of coated ECM in the COP-MPSs

Immunostaining was used to evaluate coated Matrigel or collagen type I in the COP-MPS microfluidic channels. The ECM-coated microfluidic channels were fixed with 4% (v/v) paraformaldehyde (PFA) in phosphate-buffered saline (PBS) for 20 min at 25 °C, then washed with PBS. Next, the ECM-coated channels were treated with a blocking solution of PBS containing 5% (v/v) normal goat serum, 5% (v/v) normal donkey serum, 3% (v/v) bovine serum albumin (BSA), and 0.1% (v/v) Tween-20, at 4 °C for 16 h. Then, the channels were incubated at 4 °C for 16 h with the following primary antibodies: anti-mouse laminin MEC5 (rat IgG; Catalog number ab2466; 1:50; Abcam, Cambridge, UK), anti-mouse collagen IV (rabbit IgG; 1:100; Catalog number ab19808; 1:100; Abcam, Cambridge, UK), anti-mouse entactin (Nid 1; 1:50; mouse IgG; Catalog number MBS603630; MyBioSource, San Diego, USA), or anti-rabbit collagen I (rabbit IgG; 1:50; Catalog number ab34710; Abcam, Cambridge, UK) in blocking solution at 4 °C for 16 h. The channels were incubated at 25 °C for 60 min with the corresponding secondary antibodies AlexaFluor 594 donkey anti-rabbit IgG (1:1000; Jackson ImmunoResearch, West Grove, PA, USA) and Alexa Fluor 648 donkey anti-rat IgG (1:1000; Jackson ImmunoResearch, West Grove, PA, USA) in the blocking solution. The stained samples were placed on the imaging stage of a Nikon ECLIPSE Ti inverted fluorescence microscope equipped with a CFI plan fluor 10×/0.30 NA objective lens (Nikon, Tokyo, Japan), charge-coupled device (CCD) camera (ORCA-R2; Hamamatsu Photonics, Hamamatsu City, Japan), mercury lamp (Intensilight; Nikon), XYZ automated stage (Ti-S-ER motorized stage with encoders; Nikon), and filter cubes for the fluorescence channel (TRITC and CY5 HYQ; Nikon). Once the microscopy images were acquired, CellProfiler software (Broad Institute of Harvard and MIT, Version 3.1.9) was used to quantify the stained ECMs [20]. Box plots were generated using the R software (ver. 3.5.2; https://www.r-project.org/).

### 2.6. Cell culture

SH-SY5Y human neuroblastoma cells were obtained from the American Type Culture Collection. Cells were maintained in Dulbecco’s modified Eagle’s medium (DMEM) (Sigma-Aldrich, St. Louis, MO, USA) supplemented with 10% (v/v) fetal bovine serum (FBS, Cell Culture Bioscience), 2 mM L-glutamine (200 mM, Thermo Fisher Scientific, Inc., Waltham, MA, USA), and 20 mM HEPES (Fujifilm Wako, Osaka, Japan), in a humidified incubator at 37 °C with 5% (v/v) CO_2_. The cells were flushed with trypsin/EDTA (0.04%/0.03% [v/v]) solution every four days at a ratio of 1/4.

### 2.7. Cell culture in the COP-MPS

Prior to use, the MPSs were wiped with 70% (v/v) EtOH and placed under UV light in a biosafety cabinet for a 30 min. Then, as described in Section 2.5., 10 μL of ECM solution in DMEM/F12 was introduced into each microfluidic cell culture channel and kept at 4 °C for over 24 h. Excess ECM solution was then removed, and the coated channels were washed with fresh medium.

SH-SY5Ycells cultured in a dish were harvested with 1 mL of TrypLE Express at 37 °C for 5 min, then transferred to a 15 mL centrifuge tube. Then, 4 mL of cell culture medium was added to the tube, and the cells were centrifuged at 200 × g for 3 min. Once the supernatant was decanted, the cells were resuspended in 5 mL of the medium and centrifuged at 200 × g for 3 min. Next, the cells were resuspended in the medium and introduced into the MPS microfluidic channel at 2.0 × 10^6^ cells per 10 μL. Two hours after cell seeding, fresh medium was added to the microfluidic cell culture channel to remove debris. The medium was changed every 12 h until further experiments were performed.

### 2.8. Cell apoptosis assays

Annexin V staining was performed according to the manufacturer’s instructions for Alexa Fluor 594-Annexin V conjugate (Molecular Probes, Eugene, OR, USA). Briefly, after washing with annexin-binding buffer (10 mM HEPES, pH 7.4, 140 mM NaCl, 2.5 mM CaCl_2_), the cells were stained with the Alexa Fluor 594-Annexin V conjugate at 25 °C for 15 min. Following cell fixation with 4% (v/v) PFA in PBS at 25 °C for 15 min, the cells were incubated with 300 nM of Hoechst 33258 at 25 °C for 30 min.

### 2.9. Microscopic cell imaging

Samples containing cells were placed on the imaging stage of a Nikon ECLIPSE Ti inverted fluorescence microscope equipped with a CFI plan fluor 10×/0.30 NA objective lens (Nikon, Tokyo, Japan), CCD camera (ORCA-R2; Hamamatsu Photonics, Hamamatsu City, Japan), mercury lamp (Intensilight; Nikon), XYZ automated stage (Ti-S-ER motorized stage with encoders; Nikon), and filter cubes for the fluorescence channels (DAPI, GFP HYQ, and TRITC; Nikon).

### 2.10. Single-cell profiling based on microscopic images

Once the microscopy images were acquired, CellProfiler software (Broad Institute of Harvard and MIT, Version 3.1.9) was used to identify the cells using Otsu’s method [20]. The fluorescence signals from individual cells were quantified automatically. Box plots were generated using the R software (ver. 3.5.2; https://www.r-project.org/).

### 2.11. Statistical analysis

The P values were estimated using Steel–Dwass and Wilcoxon signed-rank tests and R software (ver. 3.5.2; https://www.r-project.org/).

## 3. Results

### 3.1. Fabrication of COP-MPSs

To test the effects of solvents on COP bonding in cultured cells, the commonly used solvents toluene [21], cyclohexane [22,23], and dichloromethane [24] were evaluated; henceforth, these samples are termed COP-MPS-T, COP-MPS-C, and COP-MPS-D, respectively (**Figure 1A** and **Table 1**). For comparison, a COP-MPS was also fabricated with a VUV photobonding process, termed COP-MPS-V [17] (**Figure 1B**). The VUV photobonding process requires higher microfluidic channels for the hot-press process compared with the solvent-bonding process, so two COP-MPS microfluidic structures were prepared with heights of 280 and 320 μm (RC2 and RC3, respectively) for the solvent and VUV photobonding processes, respectively. This produced microfluidic channels with a target height of 250 μm. Because solvents might cause deformation of plastic microfluidic structures, including COP, the bonding processes for each solvent was intensively optimized for each solvent to avoid deformation (**Figure 2**). Although COP-MPS-T, COP-MPS-C, and COP-MPS-V did not show any deformation after fabrication, COP-MPS-D exhibited a deformation in the microfluidic channel caused by hot pressing after exposure to dichloromethane.

**Figure 2.**
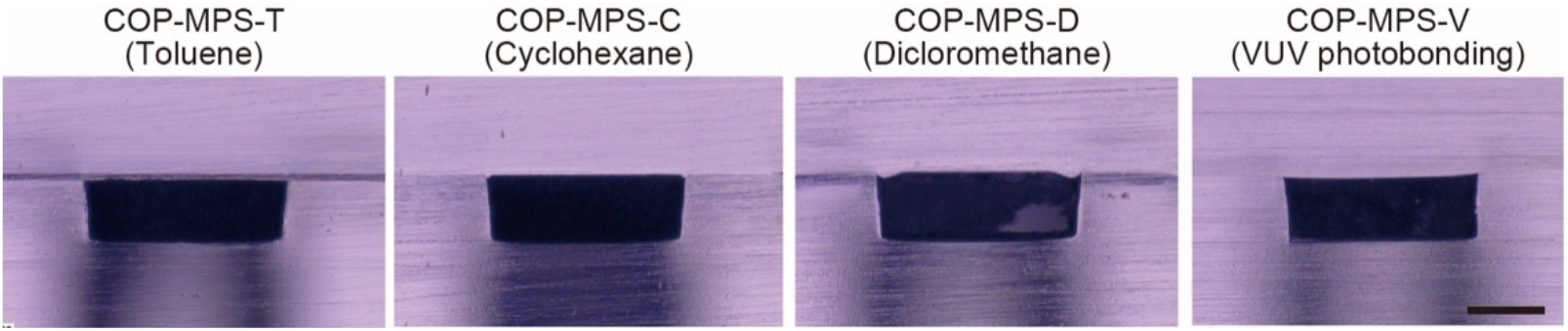
Cross sections of the COP-MPS-T, COP-MPS-C, COP-MPS-D, and COP-MPS-V specimens. Scale bar represents 300 μm.

The deformation of the COP-MPS microfluidic structures caused by the bonding processes was then evaluated (**Figure 3**). The original COP-MPS microfluidic structures (RC2 and RC3) had a coefficient of variation (CV) of less than 1% in their height and width when they were removed from the metal molds. The width of the COP-MPS-C and -V specimens decreased by 1.4% and 2.2%, respectively, in comparison to the original structures. The other specimens showed reductions of less than 1% (**Figure 3A**). VUV photobonding requires a stronger hot press for successful bonding than solvent bonding, and shrinkage of the microfluidic structure was observed in the COP-MPS-V specimen. The height of the COP-MPS-V specimen decreased by 11.2% compared to the original structure, and was close to the target height of 250 μm (**Figure 3B**). The other solvent-bonded MPSs showed reductions of approximately 3–5%, and the heights of the tested COP-MPS microfluidic channel showed a CV of 2.5%. This indicates that the solvent-bonding process did not cause deformation of the microfluidic structures. For a fair comparison of the effects of the solvents on cultured cells, it is necessary to eliminate all other factors. Hence, the effects of the solvents were investigated by further analysis.

**Figure 3.**
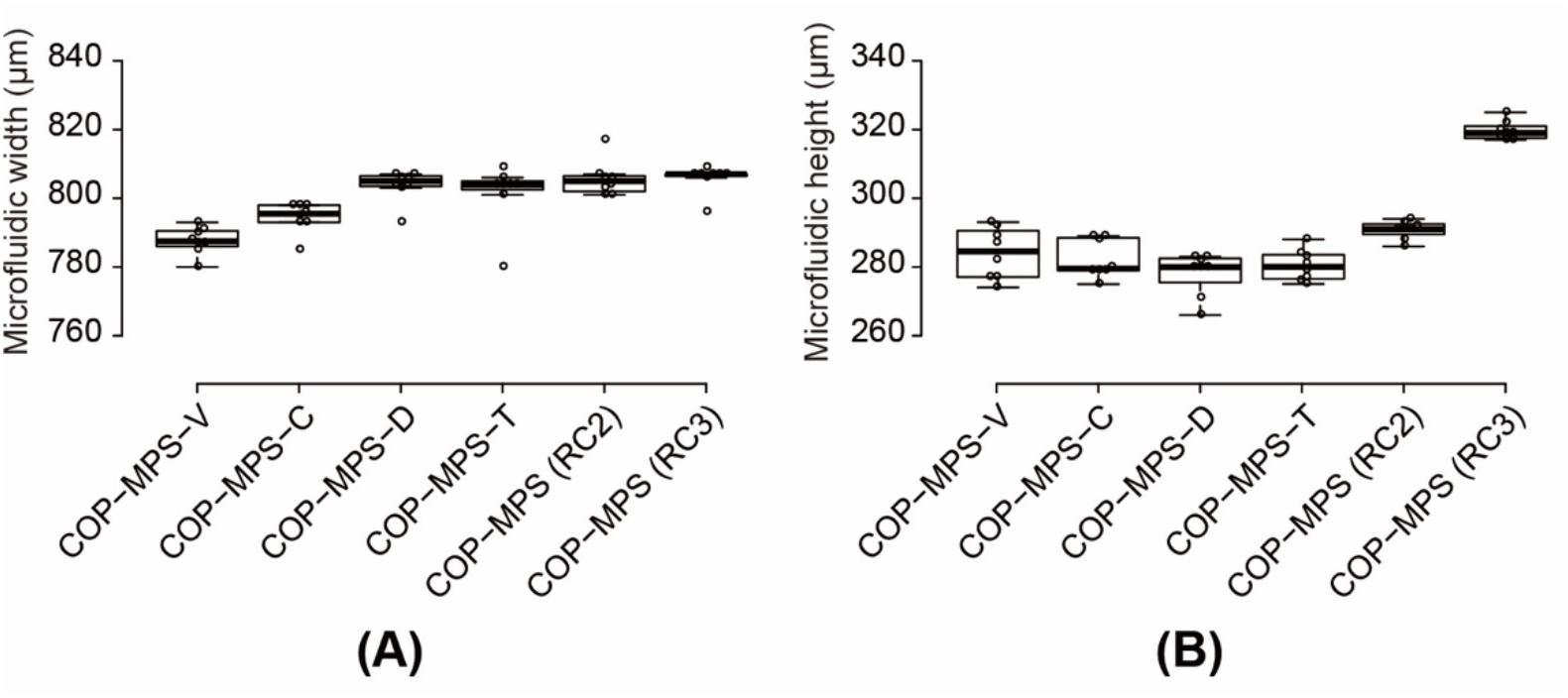
Deformation of the (A) widths and (B) heights of the VOP-MPS microfluidic channels by solvent and VUV photobonding processes. COP-MPS (RC2) and (RC3) were the original structures for solvent and VUV photobonding processes, respectively. Center lines show the medians; box limits indicate the 25th and 75th percentiles; whiskers extend to 1.5 times the interquartile range from the 25th and 75th percentiles; outliers are represented by dots; and data points are plotted as open circles. There were n = 8 sample points.

Moreover, the interface between the two assembled COP microfluidic structures in COP-MPS-T, -C, and -D exhibited color changes, as shown in **Figure 2**, but COP-MPS-V did not. This suggests that solvent bonding alters the surface properties of COP, which might affect cell cultures. In the case of COP-MPS-V, the interface could not be observed clearly, as it was tightly assembled with no obvious changes.

### 3.2. Effects of solvents on ECM coatings in COP-MPSs

To evaluate the effects of solvents on the ECM coating, immunostaining was performed for the ECM proteins on the surface of the COP microfluidic channels. There are concerns that minute solvent residues could affect the ECM coating process or disrupt the ECM proteins. Although it is difficult to distinguish between them, the effect of the residues could be evaluated by observing the coated ECMs. This study used Matrigel and collagen type I (Col I), which have been used for a variety of cell cultures, stained them with their specific antibodies, and conducted quantitative analyses. The major components of Matrigel are collagen type IV (Col IV), laminin, and entactin (also called Nidogen-1) [25–28], which were stained and analyzed with their respective antibodies.

In the case of Col IV, the COP-MPSs showed higher fluorescence in the coated microfluidic channels than in the negative control (n.c.), as shown in **Figure 4A**. Moreover, specimen COP-MPS-V had many stained clusters with strong fluorescence because of the fibrous collagen structures, which are required for the ECMs for function properly (**Figure 4B**). Due to the nature of such collagen fiber structures under *in vivo* physiological conditions, these stained clusters are a good indicator of the coating efficiency of collagens. In contrast, specimens COP-MPS-T, -C, and -D showed fewer immunostained clusters, suggesting that the solvents affected the coating process or disrupted the Matrigel coating. Moreover, the fluorescence intensity of each stained cluster had an inverse relationship with the area of the stained clusters. This suggests that the Col IV proteins attached to the COP surface treated with VUV formed larger and flatter clusters, which decreased their fluorescent intensity. Although the fluorescent intensities of the stained clusters in specimens COP-MPS-C and -T were higher than those in COP-MPS-V and COP-MPS-D, the numbers of stained clusters and the stained areas in COP-MPS-C and COP-MPS-T were much lower than those in COP-MPS-V and COP-MPS-D. This suggest that cyclohexane and toluene affected the Col IV coating.

**Figure 4.**
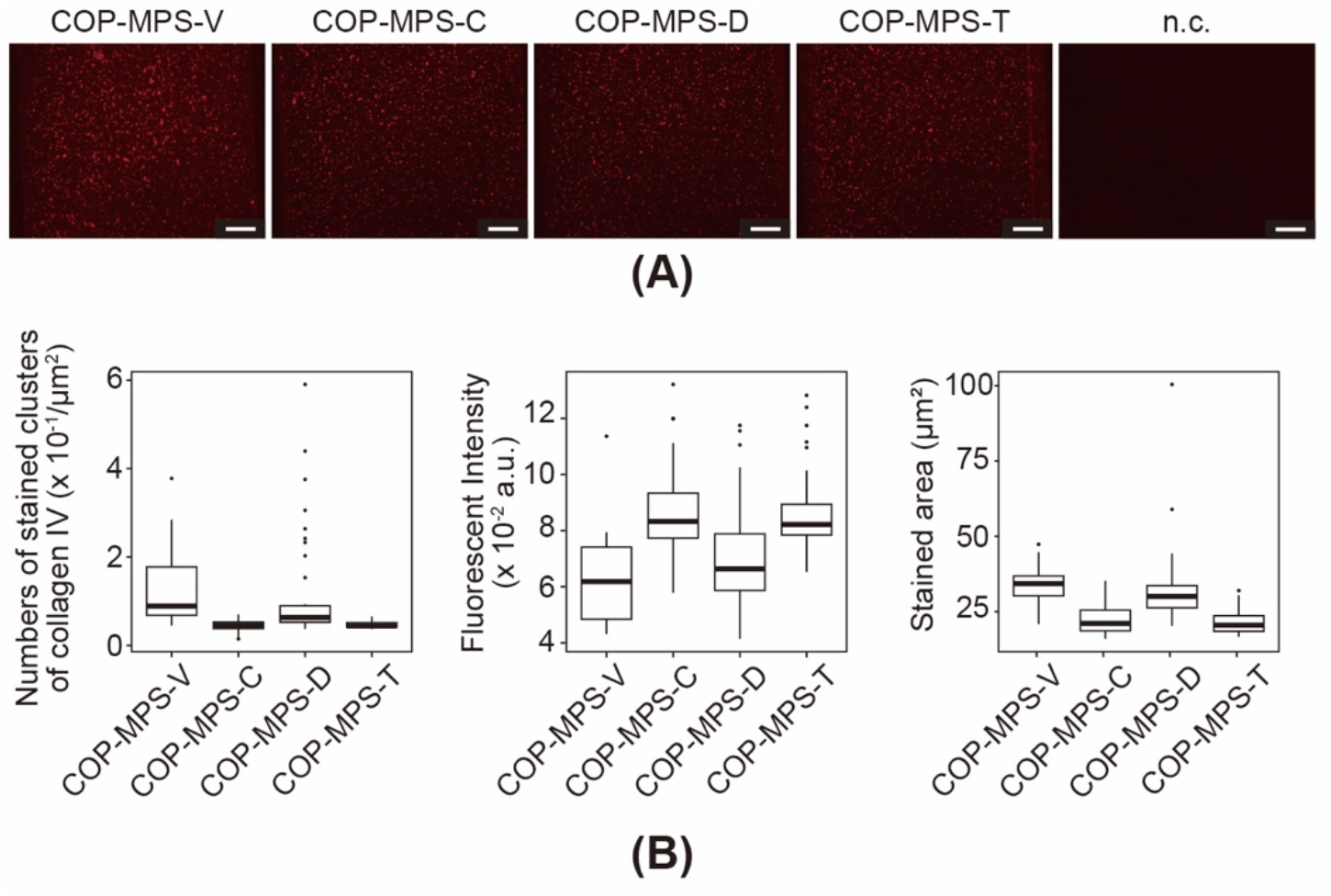
Immunofluorescent analyses of collagen IV (Col IV) in Matrigel coated COP-MPS microfluidic channels. (A) Micrographs of immunostained Col IV coated COP-MPS microfluidic channels produced using solvents or VUV photobonding processes. n.c. represents the negative control of a COP-MPS-V microfluidic channel coated with Matrigel, and only stained with a secondary antibody. Scale bars represent 100 μm. (B) Quantitative analyses of the immunostained Col IV in the coated COP-MPS microfluidic channels shown in (A). Center lines show the medians; box limits indicate the 25th and 75th percentiles; whiskers extend 1.5 times the interquartile range from the 25th and 75th percentiles; and outliers are represented by dots. Experiments were repeated four times, and more than ten images of COM-MPSs from each experiment were used for quantification.

An immunoassay for entactin in Matrigel was also performed, which confirmed that the stained microfluidic channels fluoresced (**Figure 5A**). After coating, increased fluorescence of entactin was observed in all the stained COP-MPSs compared to the negative control; in particular, specimens COP-MPS-V and -D showed stronger fluorescence. However, unlike the Col IV staining, stained clusters could not be observed. Although entactin has a domain that can bind Col IV to form the basement membrane *in vivo* [28], entactin showed less Col IV-like clusters, and covered the entirety of the microfluidic channels under *in vitro* conditions (**Figure 5B**). This is because the entactin protein itself did not have the ability to form fibrous structures like collagen. The fluorescent microscopic images in **Figure 5A** show that cyclohexane and toluene affected the entactin coating.

**Figure 5.**
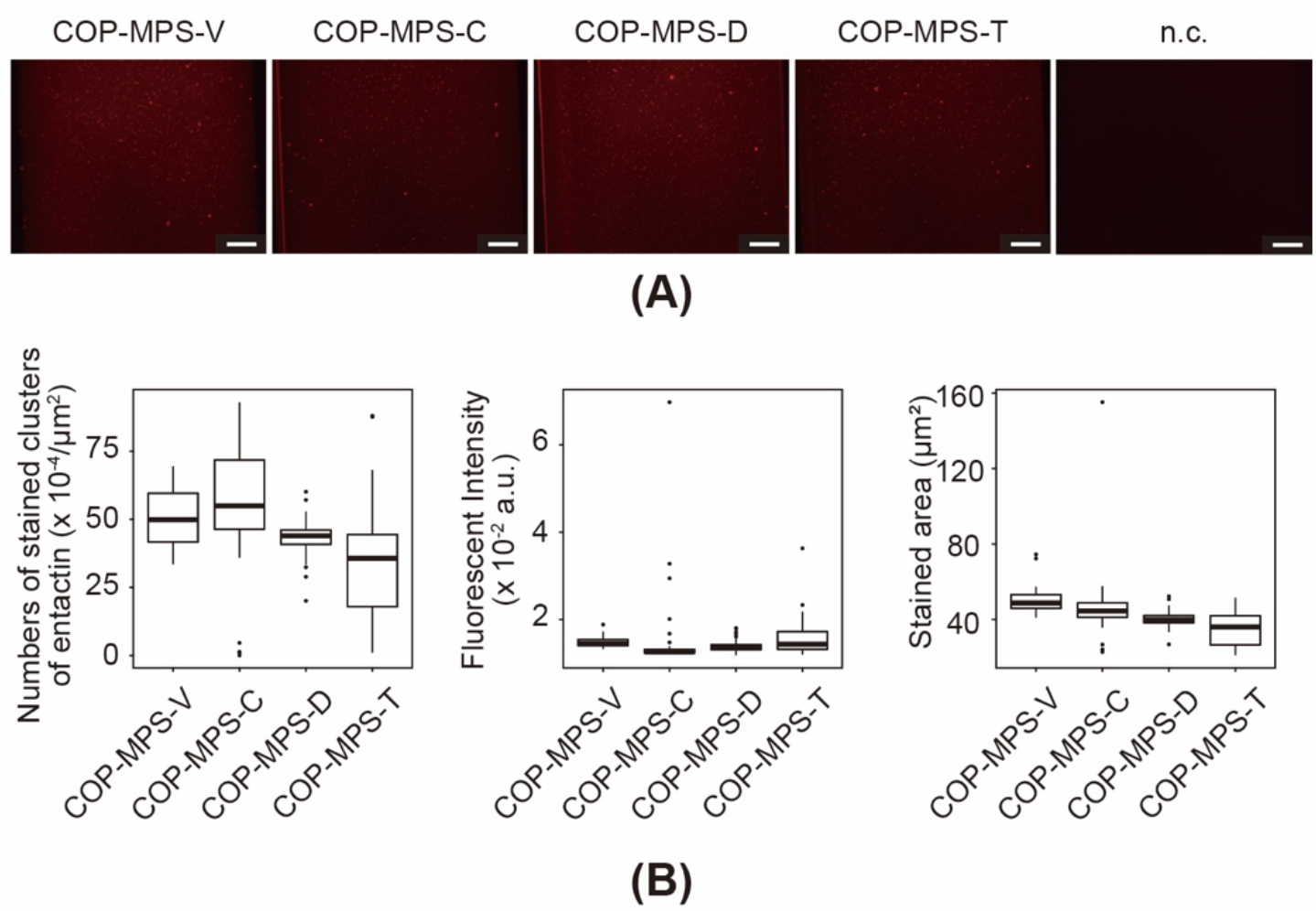
Immunofluorescent analyses of entactin in Matigel coated COP-MPS microfluidic channels. (A) Micrographs of immunostained entactin coated COP-MPS microfluidic channels produced using solvent or VUV photobonding processes. n.c. represents the negative control of a COP-MPS-V microfluidic channel coated with Matrigel, and only stained with a secondary antibody. Scale bars represent 100 μm. (B) Quantitative analyses of the immunostained entactin in the COP-MPS microfluidic channels shown in (A). Center lines show the medians; box limits indicate the 25th and 75th percentiles; whiskers extend 1.5 times the interquartile range from the 25th and 75th percentiles; and outliers are represented by dots. Experiments were repeated four times, and more than ten images of COM-MPSs from each experiment were used for quantification.

In the case of laminin in Matrigel, fluorescent signals were observed from the coated COP-MPS microfluidic channels (**Figure 6A**), and stained clusters were also observed (**Figure 6B**). Specimen COP-MPS-V showed significantly more clusters than the other COP-MPSs. Moreover, similar to Col IV, the fluorescent intensities of the stained clusters showed an inverse correlation with their area. Laminin did not form fibrous structures, but it had a strong interaction with Col IV, hence fluorescent clusters were observed. As shown in **Figure 6B**, the solvent used did not affect the number, fluorescent intensity, or stained area of the clusters, but the specimens were significantly different from COP-MPS-V.

**Figure 6.**
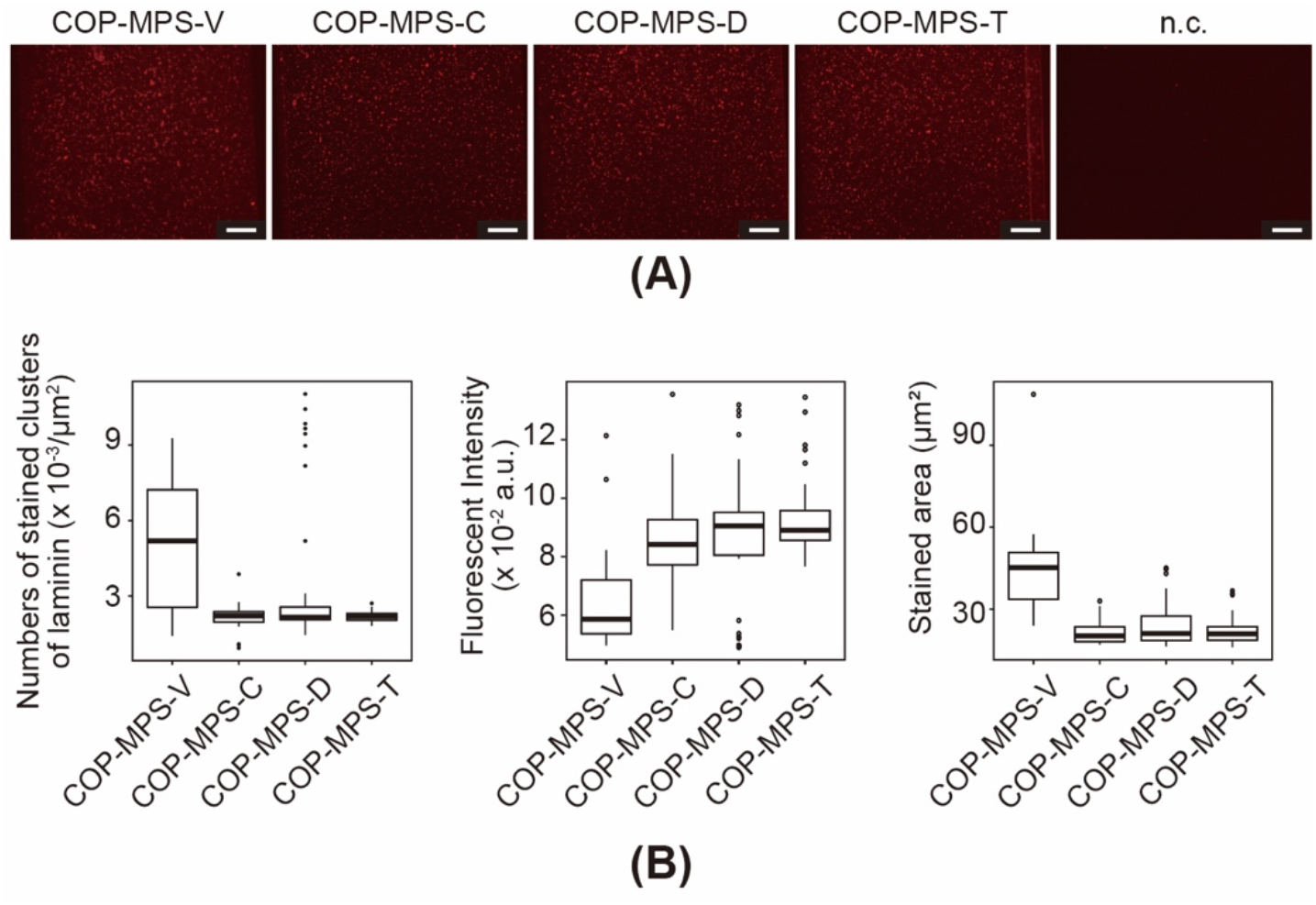
Immunofluorescent analyses for laminin in coated COP-MPS microfluidic channels. (A) Micrographs of immunostained laminin coated COP-MPS microfluidic channels produced using solvent or VUV photobonding processes. n.c. represents the negative control of a COP-MPS-V microfluidic channel coated with Matrigel, and only stained with a secondary antibody. Scale bars represent 100 μm. (B) Quantitative analyses of immunostained laminin in the COP-MPS microfluidic channels shown in (A). Center lines show the medians; box limits indicate the 25th and 75th percentiles; whiskers extend 1.5 times the interquartile range from the 25th and 75th percentiles; and outliers are represented by dots. Experiments were repeated four times, and more than ten images of COM-MPSs from each experiment were used for quantification.

In combination, this shows that the solvents have different effects on each component of Matrigel, which might affect cultured cells.

In the case with a single coating of Col I, the results were similar to those obtained for Col IV in Matrigel (**Figure 7A**). Specimen COP-MPS-V showed widely distributed fluorescent signals covering the microfluidic channels, with many fluorescent stained clusters of Col I fibers. As described, because collagen fibers are required to express collagen functionality, the presence of Col I clusters is a good indicator of coating efficiency. In comparison, specimens COP-MPS-T, -C, and -D showed fewer immunostained clusters. Quantitative analysis based on the images in **Figure 7A** was used to evaluate the Col I coating for fluorescent signals of stained Col I, as well as the number of stained clusters and their fluorescent intensities (**Figure 7B**). There were less stained clusters in specimens COP-MPS-C and -D than in COP-MPS-V and COP-MPS-T. In addition, the stained clusters in COP-MPS-V had the strongest fluorescent intensities. Interestingly, COP-MPS-V showed the least variation in the stained area, while COP-MPS-D showed the most. This suggests that the solvents altered the surface properties and interfered with the efficacy of the Col I coating.

**Figure 7.**
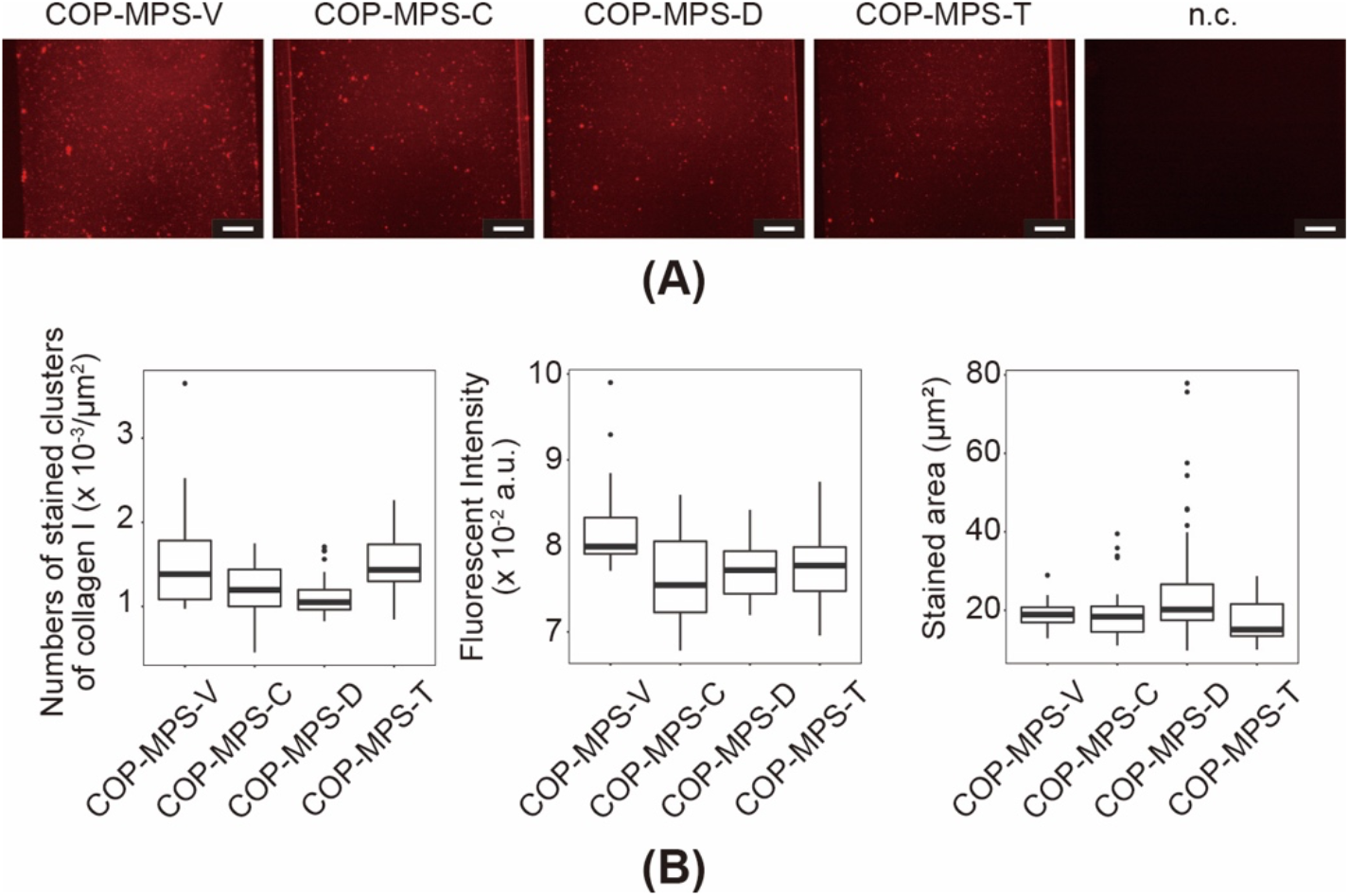
Immunofluorescent analyses for collagen I coated COP-MPS microfluidic channels. (A) Micrographs of immunostained collagen I coated COP-MPS microfluidic channels produced using solvent or VUV photobonding processes. n.c. represents the negative control of a COP-MPS-V microfluidic channel coated with Col I, and stained with only secondary antibody. Scale bars represent 100 μm. (B) Quantitative analyses of immunostained Col I in the COP-MPS microfluidic channels shown in (A). Center lines show the medians; box limits indicate the 25th and 75th percentiles; whiskers extend 1.5 times the interquartile range from the 25th and 75th percentiles; and outliers are represented by dots. Experiments were repeated four times, and more than ten images of COM-MPSs from each experiment were used for quantification.

### 3.4. Effects of solvents on SH-SY5Y cells cultured in COP-MPSs

SH-SY5Y neuroblastoma cells were used to evaluate the effects of the solvents used during manufacturing on cells cultured in COP-MPSs. Immediately after the cells were loaded into the microfluidic channels of the COP-MPSs (day 0), the SH-SY5Y cells in the Matrigel-coated COP-MPSs had a flattened cellular shape. In contrast, the cells in the Col I-coated COP-MPSs had a round cellular shape. This indicates that the cells in the Matrigel-coated COP-MPSs adhered to the substrate more efficiently (**Figure 8**). Moreover, one day after cell loading, cells in the Matrigel-coated COP-MPSs were able to adhere and spread out, but cells in the Col I-coated COP-MPSs, except COP-MPS-V, were removed by the medium change. As shown in **Figure 7**, the microfluidic channels in the COP-MPSs could be coated with Col I, but the solvents affected the coating efficiency. Consequently, the cultured cells in the Col I-coated COP-MPS could not be maintained, even for one day. This suggests that solvents might affect not only the Col I coating efficiency, but also cell adhesion or survival directly. In contrast, in the case where Col IV in Matrigel was used, the cells were maintained. This suggests that the other ECMs in the Matrigel (entactin and laminin) cooperated to support cell adhesion and survival.

**Figure 8.**
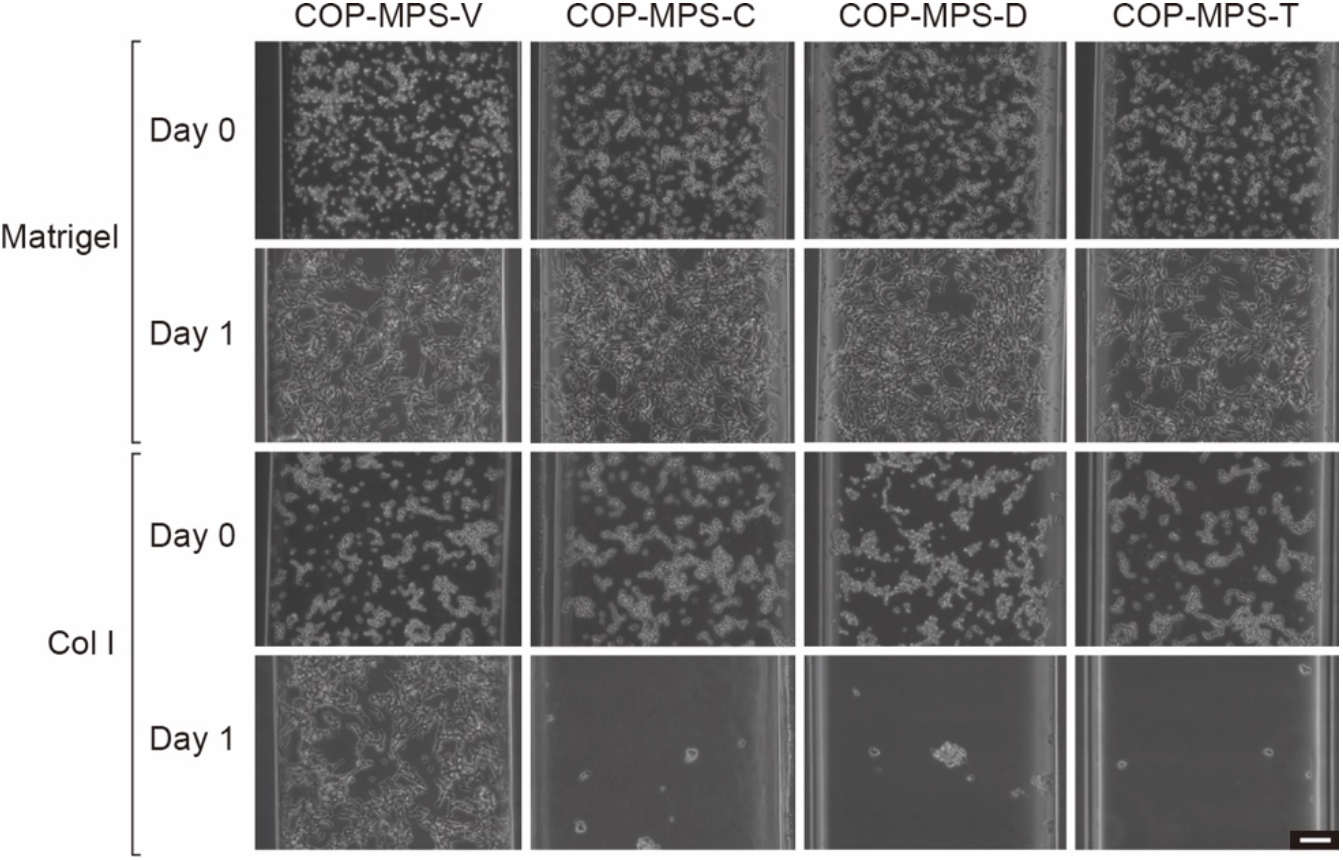
Micrographs of SH-SY5Y neuroblastoma cells cultured in COP-MPSs produced using solvent or VUV photobonding processes. The COP-MPS microfluidic channels were coated with Matrigel or collagen I (Col I). Scale bars represent 100 μm.

In the subsequent evaluations, only Matrigel-coated COP-MPSs were used.

After four days in the COP-MPSs, the cell populations and apoptotic status to the SH-SY5Y cells was evaluated by staining them with Hoechst 33258 fluorescent dye and an Annexin V apoptotic marker labeled with Alexa594 fluorescent dye (Annexin V-Alexa594) (**Figure 9A**). This was followed by quantitative single-cell profiling of the fluorescent intensity of each cell to estimate the cell densities (**Figure 9B**) and apoptotic cell numbers (**Figure 9C and Supplementary Figure S2**). The cell densities in specimen COP-MPS-V were significantly higher than those in specimens COP-MPS-C, -D, and -T (*p* = 2.4 × 10^-2^, 5.1 × 10^-4^, and 7.4 × 10^-6^, estimated using the Wilcoxon signed rank test, respectively). From the apoptosis analysis, the micrographs of cells stained with Annexin V-Alexa594 indicated that there were more positive cells in COP-MPS-C and -T than in COP-MPS-V and -D. By measuring the fluorescent intensities of the Annexin V-Alexa594 dye, apoptotic cells were counted. If the stained cells exceeded 0.1 times the fluorescent intensity of the Annexin V-Alexa594 dye they were defined as apoptosis “positive” cells (**Figure 9C**). Cells cultured in COP-MPS-V showed 4.9 ± 0.7% apoptotic cells, but those in COP-MPS-C, -D, and -T showed significantly higher numbers of 9.1 ± 2.9%, 6.7 ± 0.9%, and 7.5 ± 1.4% (mean ± S.D.; p < 0.05 compared with COP-MPS-V, estimated by Student’s *t*-test), respectively.

**Figure 9.**
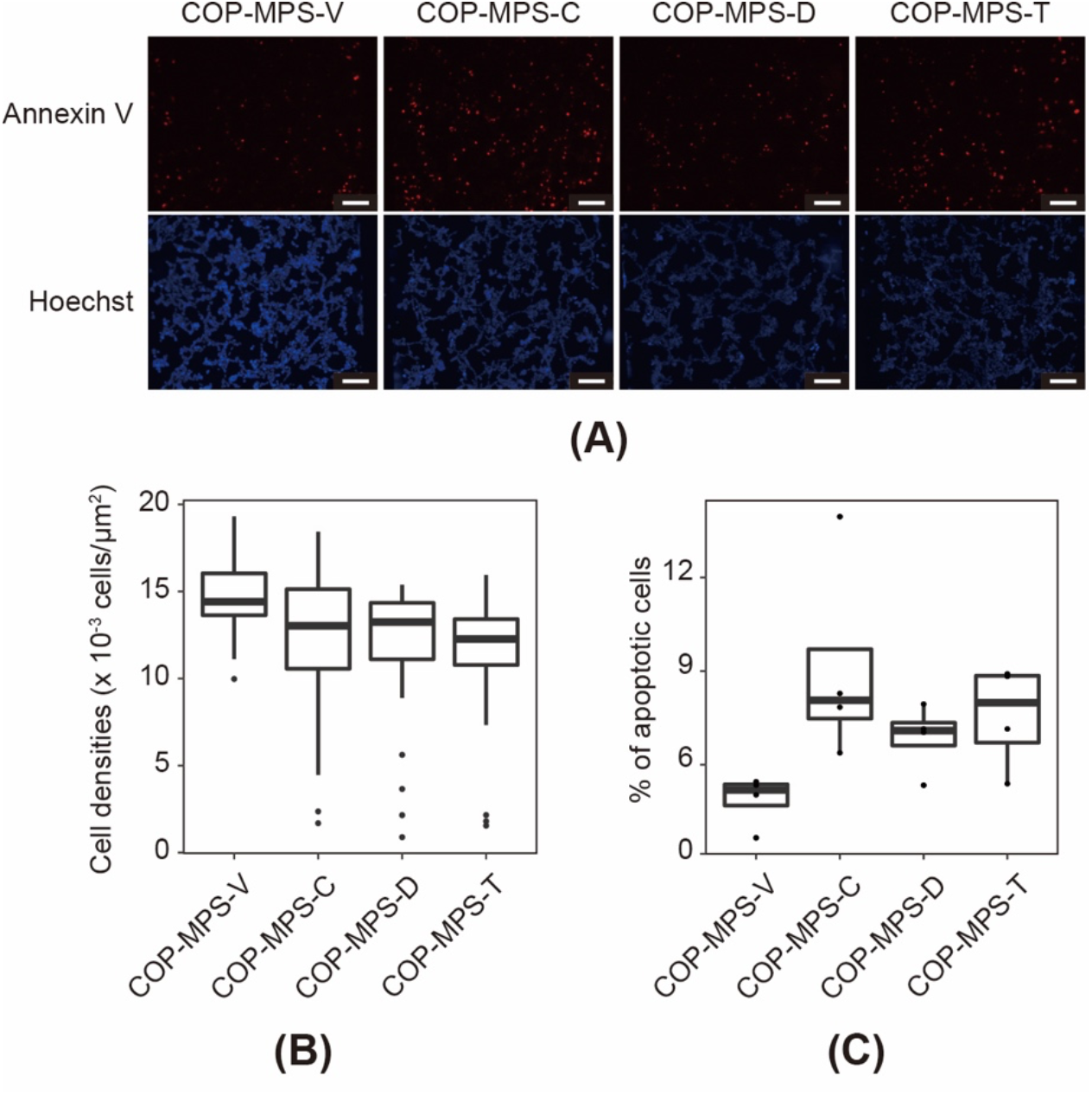
Evaluation of apoptotic cells cultured in COP-MPSs. (A) Fluorescent micrographs of SH-SY5Y neuroblastoma cells stained with Annexin V-Alexa 594 apoptotic marker. SH-SY5Y cells were cultured in Matrigel-coated COP-MPSs for four days, then stained with fluorescent dyes. Hoechst 33258 (Hoechst) fluorescent dye was used to visualize the cell nuclei. Scale bars represent 100 μm. (B) Box plots show the cellular densities after four days of culture in COP-MPSs, determined by counting the nuclei. (C) Box plots show the percentage of apoptotic cells after four days of culture in COP-MPSs determined according to the Annexin V-Alexa 594 positive cells (**Supplementary Figure S2**). Center lines show the medians; box limits indicate the 25th and 75th percentiles; whiskers extend 1.5 times the interquartile range from the 25th and 75th percentiles; and outliers are represented by dots. Experiments were repeated four times, and more than ten images of COM-MPSs from each experiment were used for quantification.

Therefore, the solvents not only disrupted the ECM coatings, but also altered the cellular status by reducing cell adhesion and inducing cellular apoptosis.

## 4. Discussion

MPSs and OoC hold great promise for the advancement of drug screening and toxicological testing, and as alternatives to animal testing, but a number of issues remain to be solved. Due to the problems faced by PDMS, most MPS researchers have focused on finding alternative microfluidic structural material [6–9]. PS, PMMA, COP, and photoresists have been used for MPS [10–14], but they might also alter cellular behaviors, and we have previously found that such microfabrication materials, including PDMS, cause changes in cellular phenotypes with changes in gene expres-sion [29]. In addition to the MPS materials, this study shows that the solvents used for bonding processes have a strong effect on cells cultured in MPSs. In fact, each material requires different bonding processes [30], so the effects of both materials and solvents on cultured cells should be investigated to prevent mistakes in the interpretation of the results.

This and our previous study [29] demonstrated an approach to assessing the materials used for MPS fabrication. This includes four key factors: (1) a microfluidic structure, (2) residue of solvents, (3) extracellular matrices, and (4) cell types, which have not been discussed in detail with respect to evaluating MPSs.

In terms of microfluidic structures, to understand these effects it is necessary to use chips with the same simple structure, but different materials and solvents, to eliminate additional parameters. This is a major challenge for MPS research. Although a number of MPSs have been established, they use different structures, materials, cell handling, and analyses. Therefore, MPS researchers and users have difficulty judging which would be best suited to their needs. Con-sequently, it is not possible to investigate and compare the effects of both materials and solvents on cultured cells in different MPSs. This and our previous study [17] used identical simple microfluidic structures with one inlet and outlet via microfluidic channels, but different materials and solvents, to provide a clear and fair comparison. This approach will help both MPS researchers and users to understand the characteristics of MPSs in detail.

Quantitative immunostaining analysis was performed to investigate whether the solvents disrupted and/or inhibited the ECM coating process. When cells are cultured in MPSs, the cell-ECM interaction is the first step, and if it does not work properly, cells cannot adhere, grow, and survive, nor be used for further cell assays. This study tested Matrigel (a mixture of Col IV, entactin, and laminin) and Col I, which are ECMs commonly used for cell cultures and MPSs. It demonstrated that the amounts and structures of the ECMs in solvent-bonded MPSs were significantly reduced. The fibrous structure of collagen means that, in addition to the fluorescent intensities, the number and area of fibrotic clusters can be used to investigate the ECM coating efficiency. Thus, it was confirmed that solvents reduced the amount of ECMs on COP microfluidic surfaces. Moreover, it is important to note that the ratios of the amounts of ECMs were also affected by solvents. Since ECM proteins trigger intracellular signaling pathways via corresponding ECM receptors (e.g., integrins) [31–34], the tendency of ECM-associated pathways would be altered by changes in the ECM ratios. Thus, it is necessary to understand the condition of coated ECMs for further cell cultures and assays.

Finally, SH-SY5Y neuroblastoma cells were used to investigate the cytotoxic effects of the solvents. By using a commonly used cell line for standardized cell-based toxicological assays [18,19], it was possible to compare the results with those from other platforms for cell cultures and assays. In this study, cell adhesion and apoptosis caused by leaked solvents were evaluated using this cell line. All the tested solvents significantly reduced cell adhesion and increased the number of apoptotic cells. As previously mentioned, the ECMs on the COP were disrupted by leaked solvents, so the results obtained for the cells should be considered with the ECM results.

This work showed an approach for evaluating the effects of solvents and materials used for MPS fabrication on cultured cells. Only SH-SY5Y neuroblastoma cells were used, but the cell type selection is important and depends on the application objectives. Since each MPS has its own objective in terms of recapitulating tissue/organ functions *in vitro*, it is necessary to select representative tissue cell lines to confirm whether they can express their proper functions in MPSs. Therefore, careful consideration should be given to the selection of cellular functional assays for evaluation. Pluripotent stem cells (PSCs) [35–37] are beneficial in this aim, because they provide differentiating cells during early development and are very sensitive to changes in their environments [38,39]. PSCs also provide a collection of tissue cells from the same cell source, so there are no concerns regarding genomic differences among established cell lines. It is also possible to evaluate the genomic abnormalities and differences in sensitivity caused by solvents and materials among tissue cells derived from PSCs. Recent advancements in a variety of omics technologies will provide deeper insights into the molecular mechanisms underlying the effects of MPS materials on cellular behaviors [40]. This information will make it possible to minimize misleading results obtained by MPSs, and to improve materials and fabrication processes to reduce undesirable effects.

MPSs can also be used for *in vitro* cancer diagnosis with very small amounts of cancer tissues from patient biopsy specimens, and to predict the efficacy and side effects of anti-cancer drugs [41–43]. Although such cancer cells seem tough with respect to their environment, these cancer specimens contain a mixture of cells and the cellular population ratios would be critical in understanding the cancer and patient characteristics. As shown, solvents and materials could change the ECMs and cultured cells, resulting in misleading outcomes for anti-cancer drug treatments.

Overall, this work presents a way to evaluate the solvents used for MPS fabrication and their effect on cultured cells. Although a number of trials have been conducted to eliminate solvents from MPSs, it is difficult to remove them completely. Such residues affected the ECM coating process and disrupted the ECM. Moreover, it was shown that SH-SY5Y neuroblastoma cells experiences apoptosis in solvent-treated MPSs.

## 5. Conclusions

This study investigated the way in which solvents used in MPS fabrication can affect cultured cells by using MPSs produced using a solvent-free VUV photobonding process for comparison. By using identical COP-MPSs for this comparison, it was possible to achieve a fair and clear assessment of the effects of solvents on cultured cells. Solvent-bonding processes can cause deformations in the microfluidic structures, but this can be minimized by optimizing the process for each solvent, and it was possible to eliminate concerns regarding inconsistencies between them. These COP-MPSs were used to investigate whether solvents affected the cell cultures, and to determine the changes in ECM efficiencies on COP surfaces. This showed that the solvents reduced the ECMs on the surfaces. Finally, the effects on cell adhesion and apoptosis, which are fundamental behaviors in cell cultures, were investigated.

The results indicated that solvent residues effected all the tested cell behaviors, resulting in improper cell cultures and assays. Thus, MPSs with solvent bonding would not be suitable for further cell cultures and assays, since the cells could not express their proper cellular status during culture. This approach for investigating the effects of the materials used for MPS fabrication is essential prior to further applications, such as drug screening and toxicological tests. Moreover, it could be the first step towards a deeper understanding of MPS components to develop MPSs with proper cellular behaviors and to advance drug screening techniques.

## Supporting information

Supplemental informations

## Supplementary Materials

The following materials are available online at www.mdpi.com/xxx/s1, Figure S1: title, Table S1: title, Video S1: title.

## Author Contributions

Conceptualization, S.T., K.H., and K.K.; methodology, X.W., S.T., and K.K.; investigation, X.W. and S.T.; resources, H.K. and K.K.; data curation, X.W. and S.T.; writing–original draft preparation, S.T. and K.K.; writing–review and editing, X.W., S.T., H.K., and K.K.; supervision, K.K.; project administration, K.H. and K.K.; funding acquisition, K.H. and K.K. All authors have read and agreed to the published version of the manuscript.

## Acknowledgments

The authors thank Shiho Terada, Satoshi Imamura, and other laboratory members for their help and valuable input. We would like to thank Editage (www.editage.com) for English language editing.

