## Supplemental informations for "Evaluation of solvents used for fabrication of microphysiological systems"

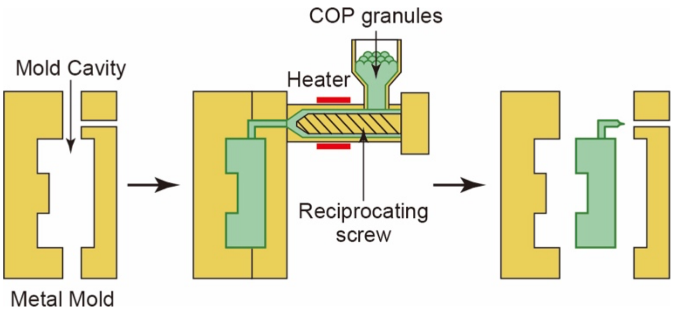


**Supplementary Figure S1. Metal molding process to fabricate the microfluidic structure of COP-MPS.**

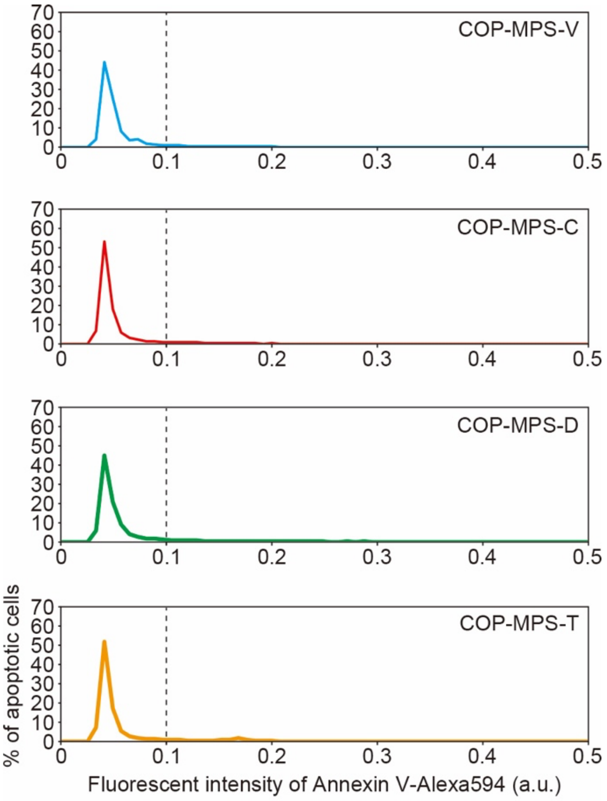


**Supplementary Figure S2. Histograms of quantitative single-cell profiling of apoptotic cells stained with Annexin V labelled with Alexa 594 fluorescent dye.** Stained cells reached over 0.1 of fluorescent intensities of Annexin V-Alexa594 dye were defined as apoptosis "positive" cells.
